# Design of Novel Selective Serotonin Reuptake Inhibitors Using Computational Modeling Studies

**DOI:** 10.1101/2022.10.10.511679

**Authors:** Lena Elisabeth H. Svanholm, Ian S. Torrence, Tram Q. Nguyen, Justin B. Siegel

## Abstract

Low levels of serotonin in the human brain have been associated with a variety of disorders including depression and anxiety. Selective serotonin reuptake inhibitors (SSRIs) are currently being prescribed to treat such conditions and examples of already marketed drugs include paroxetine, fluvoxamine, sertraline, and citalopram. Nevertheless, side effects such as nausea and drowsiness have been reported for these pharmaceuticals emphasizing the need for continuous development of new and improved lead molecules. In this study, chemical intuition and computational modeling were employed to propose two novel SSRI drug candidates with higher binding affinities to the ts3 human serotonin transporter (hSERT) than currently known SSRIs. Lastly, a homology analysis determined that *Macaca fascicularis* is a suitable model organism for future preclinical studies.

## INTRODUCTION

Selective Serotonin Reuptake Inhibitors (SSRIs) represent a class of drugs that can be used to treat various disorders including obsessive-compulsive disorder, depression, and anxiety.^1^ Depression and anxiety are widespread, debilitating disorders that affect about one in five people of clinical severity, and common symptoms include loss of pleasure, disrupted sleeping, and decreased memory.^2^ The causes underlying these symptoms are neuronal atrophy and reduced synaptic density, however, SSRIs can help alleviate these effects by locally increasing the concentration of serotonin neurotransmitters in the synaptic cleft. Specifically, the primary biological target of these inhibitors is the human serotonin transporter (hSERT) which utilizes energetically favorable cotransport of sodium and chloride ions to remove serotonin from the extracellular space and transport it across membranes.^3–5^ Serotonin itself, or 5-hydroxytryptamine (5-HT), is a neurotransmitter and a hormone that regulates various physiological cycles and responses including the sleep/wake cycle, thermoregulation, hunger, mood, digestion, memory, and pain.^6^ Usually, 5-HT is released from presynaptic neurons in the central nervous system and binds to serotonin receptors. However, studies have shown that SSRIs can bind with high affinity and specificity to the central substrate-binding site of the hSERT.^7^ This binding inhibits the reuptake of 5-HT into the presynaptic neurons and causes the protein to adopt an outward-open conformation which blocks protein activity.^7,8^

In this way, SSRIs can be classified as a class of competitive drug inhibitors and the hSERT is a popular drug target for such antidepressants that help treat both affective and neurological disorders. Yet, side effects reported for SSRIs include nausea, drowsiness, and delayed onset of antidepressant response^1^ which emphasizes the need for continuous drug optimization to develop new lead candidates that potentially result in fewer side effects. In this study, two novel SSRI candidates were designed using either algorithmically proposed bioisosteres or chemical intuition guided by computational evaluation of the predicted protein-ligand interface. The candidates were devised based on the structures of known SSRIs and on a pharmacophore model of the hSERT receptor. Figure 1 illustrates the chemical structures of these compounds.

**Figure 1.**
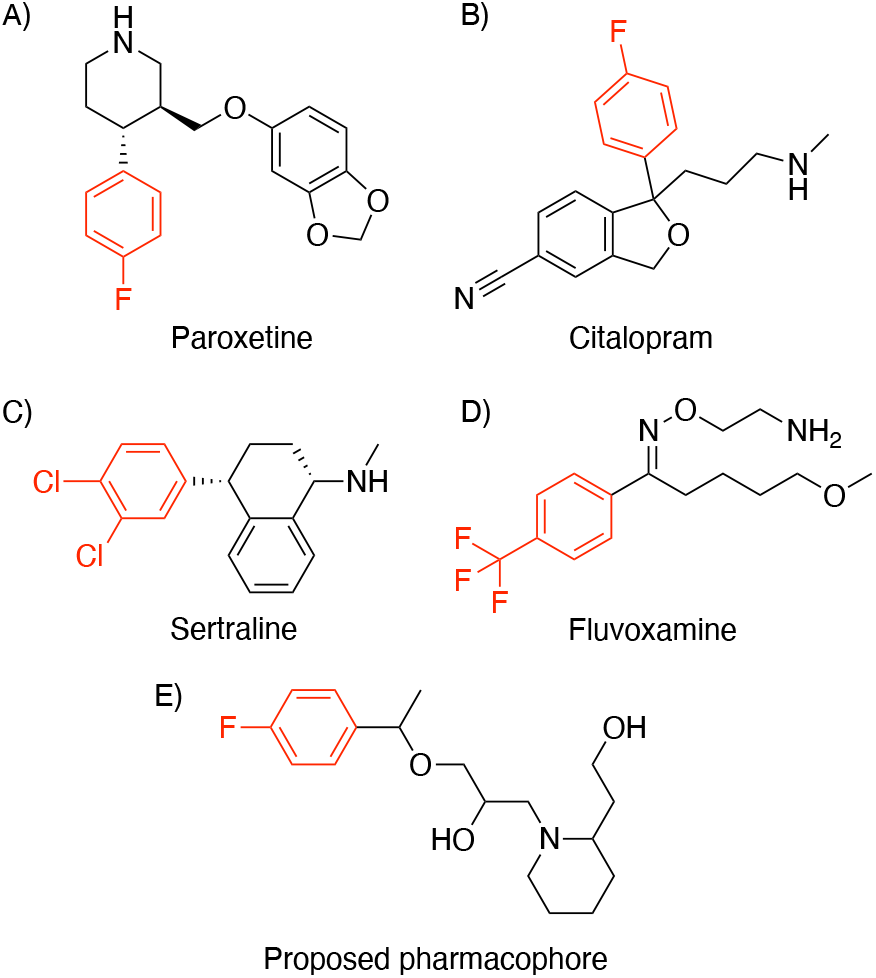
Chemical structure of A) Paroxetine, B) Citalopram, C) Sertraline, D) Fluvoxamine, and E) A proposed pharmacophore of the hSERT receptor.^3^ Structural pharmacophore elements that are retained in all SSRIs are highlighted in red directly on the figure.

This specific pharmacophore was proposed after virtual screening of a small molecule library of a series of hSERT inhibitors^3^ and the model served as an initial guideline for SSRI design in this study. In addition, docking calculations were performed on both candidates to help predict each molecule’s affinity for the receptor. Lastly, homology analyses were conducted to identify any potential off-target effects as well as to identify an organism suitable for clinical trials.

## METHODS

The crystal structure of the ts3 human serotonin transporter complexed with sertraline at the central site (PDB ID: 6AWO)^9^ was obtained from the Protein Data Bank and used to perform docking simulations of four known SSRIs (citalopram, sertraline, fluvoxamine, and paroxetine) and of developed drug candidates. First, 3D structures of the molecules were constructed using *GaussView*^10^ whereas *Gaussian 09W*^11^ was used to geometrically optimize these structures using the AM1 semiempirical approach. Next, the OpenEye software, *MakeReceptor*^12^, was used to define the active site shape of the ts3 hSERT and bioisosteres of sertraline were generated and ranked with *vBrood*^13^ based on shape and electrostatic environment. In addition, *OMEGA*^14^ was used to generate all possible conformations of each molecule obtained by C-H sigma bond rotations. The conformer libraries were docked into the active site of the ts3 hSERT using *FRED*^15^ and the molecules were ranked by a docking score according to their binding affinity of the proposed active site as determined by shape, hydrogen bonds, as well as protein and ligand desolvation.

Once the drug candidates were chosen, *SciFinder*^16^ was used to confirm that these were not already proposed in the literature. A property screening of the drug candidate library was then performed using OpenEye’s *FILTER*^17^ software. This molecular filter screened the compounds based on selected ADMET properties and on functional groups. Protein-ligand interactions between the drugs and the ts3 hSERT as well as bond distance measurements between catalytic site residues and the ligands were visualized in *PyMol*^18^. Then, *BLAST*^19^ was used to conduct homology analyses and search for serotonin transporter protein homologs. Finally, *JalView*^20^ was used to analyze and compare protein sequences of various model organisms.

## RESULTS AND DISCUSSION

### Evaluation of Binding Affinities and ADMET Properties of Known SSRIs

When initiating any drug development process that seeks to improve either pharmacokinetic or pharmacodynamic properties of established drugs, property screenings of the known ligands must be conducted. These screenings serve as control experiments during which structural characteristics are compared to binding properties and established ADMET predictors. In addition, such controls can help identify any areas that may serve as the focus of structural optimization. For these reasons, the 3D structures of four known SSRIs – paroxetine, citalopram, sertraline, and fluvoxamine – were first built and optimized in *GaussView*^10^ and *Gaussian 09W*^11^, respectively. A 3D model of the ts3 hSERT receptor was then constructed and the active site was defined with molecular and spatial constraints that ensured that critical residues known to form protein-ligand interactions with various SSRIs were contained within the site. Examples include Asp98, Ile172, Phe335, and Ser336 which have been reported as residues important for SSRI catalytic site ligand binding.^1,21–23^ Specifically, these residues help stabilize the protein in the active site by hydrogen-bonding, hydrophobic, and π-stacking electrostatic interactions. Figure 2 illustrates both the relative locations of these critical residues in the ts3 hSERT as well as the placement of sertraline in the catalytic pocket. The residues Asp98, Ile172, Phe335, and Ser336 are highlighted in purple in all subsequent protein-ligand interaction figures.

**Figure 2.**
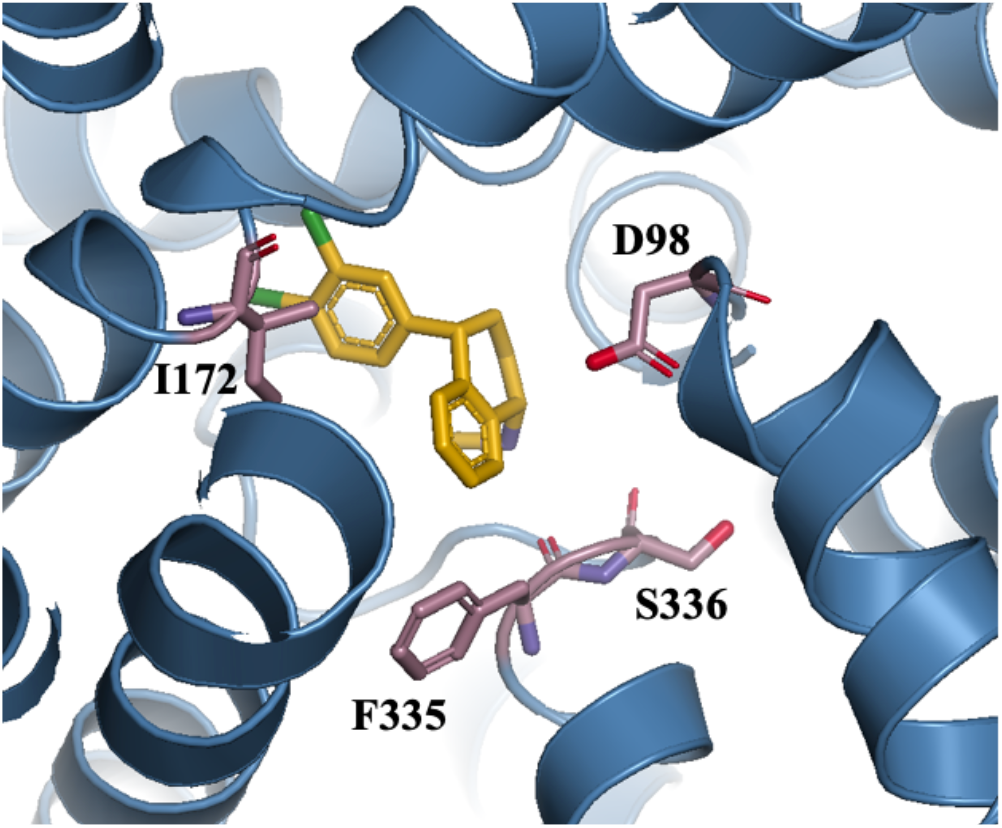
Zoomed-out view of the t3 hSERT active site complexed with the ligand sertraline (PDB ID: 6AWO)^9^ in gold. Asp98, Ile172, Phe335, and Ser336 are drawn as purple sticks.

Once this catalytic site was built, the known ligands were docked into the protein. The binding affinity of each drug with the ts3 hSERT was evaluated and quantified by docking score values as well as additional features that drugs are commonly evaluated for, reported in Table 1. The detailed docking reports and calculations are available in the Supporting Information “Docking Reports for SSRIs and Proposed Drug Candidates”. Additional structural characteristics of sertraline that contributed to the decision of making this drug the lead compound to build from in this study include the number of Lipinski hydrogen-bond acceptors as well as the number of rotatable bonds. Specifically, the H-bond acceptor count in sertraline is only one which is lower than that of both citalopram (three) and fluvoxamine (four). Increasing this count served as inspiration for the design of a new lead molecule. Lastly, the number of rotatable bonds is two for sertraline whereas this number is five for citalopram and ten for fluvoxamine. Both of these molecules had better docking scores than sertraline and since an increased number of rotatable bonds can allow a molecule to adopt different conformations in a protein binding pocket, this characteristic provided additional ideas for structural optimization.

**Table 1.** ADMET properties and selected characteristics of four known SSRIs – paroxetine, citalopram, fluvoxamine, and sertraline.

|  | Paroxetine | Citalopram | Fluvoxamine | Sertraline |
| --- | --- | --- | --- | --- |
| Docking score | -7.41 | -10.78 | -10.44 | -8.02 |
| Molecular Weight (g/mol) | 330.4 | 325.4 | 319.3 | 307.2 |
| Lipinski H-bond donors | 1 | 1 | 1 | 1 |
| Lipinski H-bond acceptors | 4 | 3 | 4 | 1 |
| XLogP | 3.37 | 4.18 | 4.07 | 5.50 |
| Chiral centers | 2 | 1 | 0 | 2 |
| Rotatable bonds | 4 | 5 | 10 | 2 |

Overall, the higher XLogP value, the lower docking score, and the low numbers of H-bond acceptors and rotatable bonds in sertraline helped identify this SSRI as the lead candidate. The study aimed to achieve two main goals; 1) create a drug candidate with no Lipinski violations and 2) create a drug candidate with a significantly improved docking score. Before this process was initiated, the protein-drug interactions formed between sertraline and the ts3 hSERT active site were illustrated in *PyMol*^18^ to understand which regions of the drug were already engaged in bonding interactions and which regions could serve as potential sites of new interactions.

Figure 3 shows both the 2D and 3D molecular structures of sertraline as well as bond distances and electrostatic interactions formed between catalytic site residues in the ts3 hSERT and the drug. The figure illustrates how sertraline is stabilized in the central binding pocket of the ts3 hSERT primarily by hydrophobic and π-stacking interactions formed from nearby residues with the tetralin ring and with the para- and meta-substituted dichlorophenyl ring.

**Figure 3.**
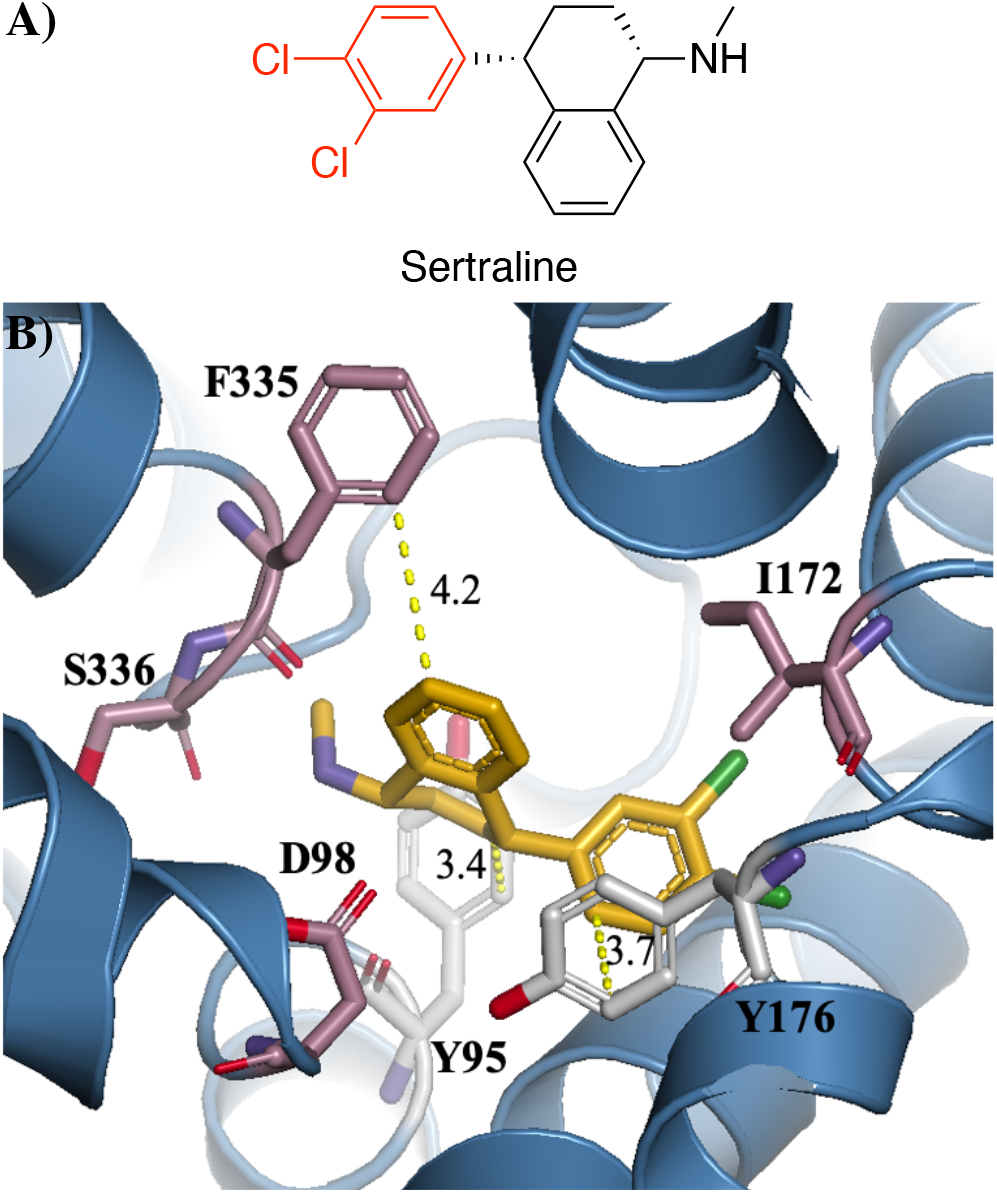
A) The 2D molecular structure of sertraline. B) Electrostatic interactions formed between sertraline and residues Y95 (hydrophobic), Y176 (π-stacking), and F335 (hydrophobic) in the catalytic site of the ts3 hSERT. Distances are reported in Ångströms (Å).

The molecule also contains a secondary amine which is capable of hydrogen bonding. However, the *FRED*^15^ report of sertraline docked into the active site of the ts3 hSERT, cf. Supporting Information “Docking Reports for SSRIs and Proposed Drug Candidates”, did not show any hydrogen-bonding interactions in the residue fingerprint section. Based on this observation, the aliphatic amine became the focus of structural optimization.

### Computationally Based Drug Design: Drug Candidate 1

Starting with a computational approach, bioisosteres of sertraline were generated using *vBrood*^13^ and a small molecule library of ten compounds was produced. These were docked into the active site of the hSERT protein. Next, the leads were scored and ranked based on their hydrogen bonding abilities with the catalytic pocket and based on how well the shape of the ligand fit into the pocket. The top hit of this search found that adding a hydrophobic cyclopropyl group with a hydrophilic hydroxyl substituent to the tetralin ring provided a docking score of -11.88 which is a 48 % improvement as compared to unmodified sertraline. The hypothesis was that this addition would enable the molecule to participate in both hydrogen-bonding and hydrophobic interactions with the active site, however, the residue fingerprint showed that the main source of the increased docking score was a hydrogen-bonding interaction alone. Specifically, the drug candidate was stabilized in the active site by an H-bond with Ser336. This gave a hydrogen-bond docking score of -1.18 rather than -0.05, cf. Table 2, illustrating that this interaction is important for binding. Another electrostatic interaction formed was π-stacking with Phe335 which is one of the residues reported as critical for catalytic site SSRI binding.^21–23^ The 2D and 3D dimensional structures of the candidate are illustrated in Figure 4A and Figure 4C, respectively.

**Table 2.** Selected ADMET properties and docking score values of sertraline, drug candidate 1, and drug candidate 2.

|  | Sertraline | Candidate 1 | Candidate 2 |
| --- | --- | --- | --- |
| Molecular Weight (g/mol) | 307.2 | 333.3 | 351.3 |
| Ratio Lipinski H-Bond Donors: Acceptors | 1: 1 | 1: 1 | 1: 2 |
| XLogP | 5.50 | 5.28 | 4.83 |
| Chiral Centers | 2 | 4 | 2 |
| Rotatable Bonds | 2 | 2 | 5 |
| 2D Polar Surface Area | 16.61 | 20.23 | 29.46 |
| H-Bond Docking Score | -0.05 | -1.18 | -1.22 |
| Shape Docking Score | -11.43 | -14.59 | -14.43 |
| Total Docking Score | -8.02 | -11.88 | -11.56 |
| Total Docking Score Improvement (%) | - | 48 % | 44 % |

**Figure 4.**
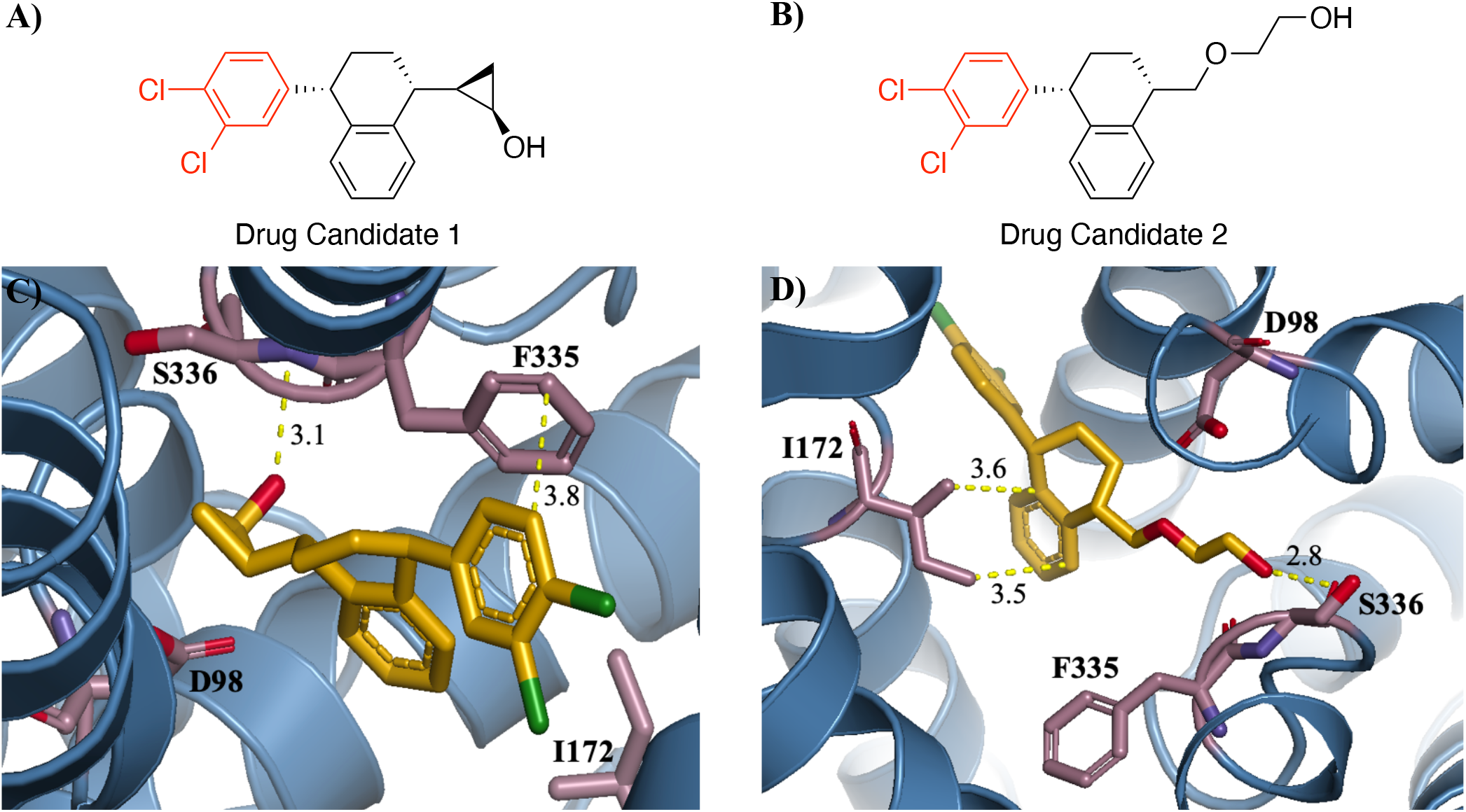
A) The 2D molecular structure of computationally designed drug candidate 1. B) The 2D molecular structure of drug candidate 2. C) Electrostatic interactions formed between drug candidate 1 and the residues F335 (π-stacking) and S336 (hydrogen-bonding) in the ts3 hSERT active site. D) Electrostatic interactions formed between drug candidate 2 and the residues I172 (hydrophobic) and S336 (hydrogen-bonding) in the ts3 hSERT active site. Distances are reported in Ångströms (Å).

The XLogP value of the proposed molecule was also calculated and the value was found to be 5.28, cf. Table 2. While this Lipinski violation does not specify that the drug cannot be orally bioavailable, it indicates that oral administration may be difficult. Specifically, highly lipophilic drugs with XLogP values above 5 can have increased molecular promiscuity and toxicity due to their ability to bind to hydrophobic targets other than the desired one(s).^24^ For this reason, an additional drug candidate was proposed using chemical intuition, guided by the structural analysis, with the purpose of obtaining an XLogP value below 5 while also maintaining the improved predicted binding affinity to the ts3 hSERT.

### Chemical Intuition Based Drug Design: Drug Candidate 2

The model was driven by the hypothesis that adding a carbon alkyl chain with a polar atom incorporated would increase the polar surface area (PSA) of the compound. Increasing the PSA was a focus of optimization since having more polar atoms can contribute to decreased lipophilicity and thus to a lower XLogP value. In addition, the model design was guided by the proposed pharmacophore in Figure 1 which includes an ether-containing 5-membered alkyl chain inserted between two ring structures. Lastly, introducing a hydroxyl group in the terminal end of the chain would expectedly allow the molecule to form strong hydrogen-bonding interactions with the protein.

Overall, these hypotheses were confirmed by docking and binding studies which showed that the ligand participated in hydrophobic interactions with Ile172 and in H-bonding interactions with Ser336 as illustrated in Figure 4D.

With these structural modifications, the overall docking score of drug candidate 2 was improved by 44 % as compared to unmodified sertraline. In addition to the newly formed hydrogen-bond, a source of improved binding was better spacefilling of the catalytic pocket as represented by an improved shape docking score. Furthermore, replacing the amine in sertraline with an alkyl chain resulted in an XLogP value below 5. Specifically, the predicted XLogP is 4.83, cf. Table 2, which could be accounted for by a doubled 2D PSA value. Comparisons of these mentioned values and of additional ADMET properties for sertraline and for the two proposed drug candidates are found in Table 2. Lastly, a chemical structure search in *SciFinder* confirmed that neither drug candidate 1 nor 2 are proposed in existing literature, cf. Fig. S-1 and Fig. S-2 in the Supporting Information “Homology Analyses of the ts3 hSERT”.

To illustrate the structural differences responsible for the changes in predicted binding modes, Figure 5 is included. This figure shows representations of sertraline and of both candidates oriented the same way in the ts3 hSERT catalytic pocket. Nearby residues are highlighted for context. The figure helps illustrate differences in catalytic site space fillings and in predicted electrostatic interactions that occur between the protein and the drug candidates. As an example, Figure 5A and 5B show that the side chain of Ile172 is located nearby the para- and meta-substituted dichlorophenyl ring in sertraline and drug candidate 1, respectively.

**Figure 5.**
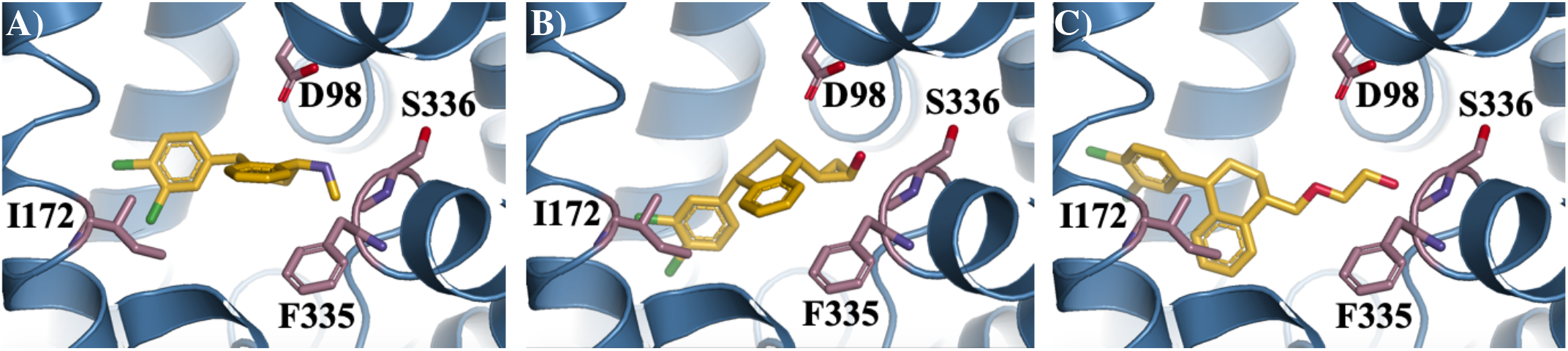
The t3 hSERT active site complexed with: A) Sertraline, B) Drug candidate 1, and C) Drug candidate 2. Selected nearby residues – Asp98, Ile172, Phe335, and Ser336 – are represented as purple sticks and oriented identically in all three schemes.

However, for drug candidate 2, the hydrophobic side chain is located near the nonpolar tetralin ring, cf. Figure 5C. This interaction predicts an increase in active site stabilization that does not occur for the other ligands. In this way, small changes in the identity of the tetralin-substituents result in changes in the structural orientation of the ligands in the ts3 hSERT active site which cause changes in predicted ADMET properties and binding modes.

In addition to an improved docking score and a lower XLogP value, the number of rotatable bonds also differs between sertraline (two) and drug candidate 2 (five) as seen in Table 2. A study investigating the structural similarities of drugs, human metabolites, and toxins found that 79 % of current drug candidates contain between one and ten rotatable bonds with a mean value of six bonds per drug.^25^ Since a rotatable bond count of five falls within that range, this is not a factor expected to cause complications. In fact, this characteristic may even contribute to the increased binding affinity. Nevertheless, the chiral center count of drug candidate 1 (four) can potentially be problematic. Specifically, having more chiral centers can make the practical isolation and purification of a drug more difficult in a chemical synthesis process and can also provide unexpected activity since the pharmacological and physiochemical properties of enantiomers may differ significantly.^26^ Furthermore, optically pure substances can undergo racemization in the human body making it necessary to map out the activity, and to conduct extensive biological testing, of all enantiomers and diastereomers.

While these concerns present limitations of drug candidate 1, a potential benefit of both proposed candidates is that the route of metabolism will be different from that of unmodified sertraline. Specifically, sertraline undergoes extensive first-pass metabolism whereby higher drug doses are needed to obtain therapeutic effects.^27^ Major metabolites are achieved by N-dealkylation and by oxidative deamination of the secondary amine by various CYP450 isoforms^28^, however, since neither proposed candidate contains an amine, alternate routes of metabolism must occur for these drugs. This hypothesis, however, was not computationally examined as there is no reliable predictor of biological responses to drugs; a current limitation in the field of computational drug design. In practice, the human body is an intricate system and many enzymes, hormones, and metabolites, as well as co-administration of other pharmaceuticals, may affect the performance of drugs and their compatibility with the body. To understand the possibilities of off-target drug effects and to identify organisms suitable for biological testing and clinical trials, a homology analysis was performed.

### Investigation of Potential Clinical Trial Organisms and Possible Off-Target Effects

First, the possibilities of off-target drug effects with both beneficial and pathogenic organisms were checked. A homology analysis of the ts3 hSERT and three beneficial microbes – *Lactobacillus, Bifidobacterium*, and *A. muciniphila* – found that the largest percent identity alignment was 27.1 % with the sodium: calcium symporter of *A. muciniphila*. Similarly, a homology analysis comparing the hSERT with four pathogenic microbes – *Staphylococcus aureus, Haemophilus influenzae, Candida albicans*, and *Tinea* – found a maximum percent identity alignment of 28.6 % with a flu microbe. This suggests that neither beneficial nor pathogenic homologs have structurally related proteins to the ts3 protein, minimizing the chance of offtarget effects of the drugs with these common microbes.

Lastly, the sequence of the ts3 hSERT was compared with those of common clinical trial organisms; mice, rabbits, beagles, and macaques. Detailed homology analyses for these organisms and for the discussed beneficial and pathogenic ones can be found in the Supporting Information. The investigation classified a macaque as a suitable clinical trial organism. Specifically, the sodium-dependent serotonin transporter in *Macaca fascicularis* shares 97.8 % sequence identity with the human counterpart. This suggests that the interactions predicted to form between the drug candidates and the hSERT can also occur in the animal model. In addition, Ile172, Phe335, and Ser336 are highly conserved residues in the homologous macaque protein further supporting the perspectives in using this organism as a model system. In fact, Ser336, which participates in hydrogen-bonding interactions responsible for improved docking scores, is conserved in all four macaque models and in 91 % of all animal models studied. Overall, since the homolog protein is also a serotonin transporter, it is likely that the protein-drug interactions formed in the macaque will resemble those in the human body emphasizing the perspectives and promises in conducting further metabolism, toxicity, and clinical studies with the proposed drug candidates.

## CONCLUSION

In this study, two novel SSRI drug candidates were proposed based on the structure of sertraline, on a pharmacophore model of the ts3 hSERT active site, on computational modeling studies, and on protein-ligand structure-guided chemical intuition. The developed leads contained no known aggregators nor toxicophores and both compounds satisfied common features sought after to increase the probability of ADMET properties for orally bioavailable drugs. In addition, candidates 1 and 2 exhibited total docking score improvements of 48 % and 44 %, respectively, as compared to unmodified sertraline. The primary sources of increased docking scores were better catalytic site pocket filling as well as hydrogen-bonding interactions formed between a hydroxyl substituent in the candidate and Ser336 in the ts3 hSERT. In addition, drug candidate 2 had an XLogP value of 4.83 thus exhibiting no Lipinski violations as opposed to both sertraline and drug candidate 1. These improved binding affinities to the ts3 hSERT active site illustrate that the proposed leads have the potential to serve as new treatment options for various disorders associated with deficient serotonin levels. Nevertheless, a limitation of the study is that it was unable to draw any conclusions on unexpected enantiomer activity and on metabolic byproducts. To establish these characteristics, biological testing is needed and a homology analysis found *Macaca fascicularis* to be a suitable model organism for clinical studies.

## Supporting information

Supplemental Docking Reports

Supplemental Homology Analyses of the ts3 hSERT

## ASSOCIATED CONTENT

### Supporting Information

The following Supporting Information is available free of charge on the bioRxiv website.

Docking Reports for SSRIs and Proposed Drug Candidates (PDF) Homology Analyses of the ts3 hSERT (PDF)

## AUTHOR INFORMATION

### Author Contributions

Research was designed by all authors; all experiments were carried out by L.C.R. The manuscript was written through contributions of all authors. All authors have given approval to the final version of the manuscript.

## ACKNOWLEDGMENT

Research reported in this publication was supported by UC Davis, the National Science Foundation Award Numbers 1827246, 1805510, 1627539, the National Institute of Environmental Health Sciences of the National Institutes of Health (NIH) under Award Number P42ES004699, UC Davis, NIH Award Number R01 GM 076324-11 and the Rosetta Commons. The content is solely the responsibility of the authors and does not necessarily represent the views of the National Institutes of Health or National Science Foundation. This study was derived from a course based undergraduate research study conducted in Chemistry 130B at UC Davis.

## ABBREVIATIONS

SSRI: selective serotonin reuptake inhibitors
hSERT: human serotonin transporter
5-HT: 5-hydroxytryptamine

