## Supplemental Docking Reports for "Design of Novel Selective Serotonin Reuptake Inhibitors Using Computational Modeling Studies"

### Supporting Information – Docking Reports for SSRIs and Proposed Drug Candidates

| | |
|---|---|
| Molecule Name | Citalopram |
| Molecular Weight | 324.4 |
| XLogP | 4.2 |
| PSA | 36.3 |
| Heavy Atoms | 24 |
| Acceptor Count | 3 |
| Donor Count | 0 |
| Chelator Count | 0 |

### Total Score -10.78

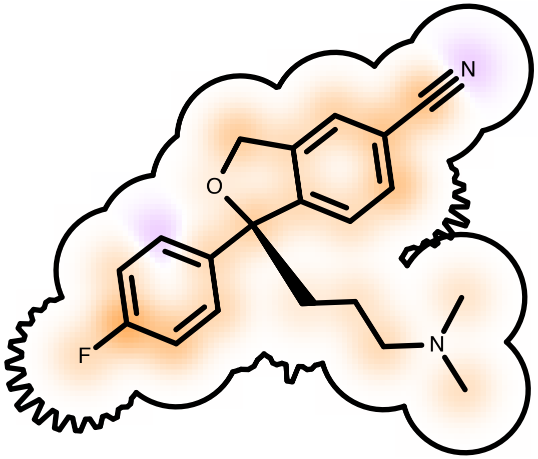

Score compared to other molecules

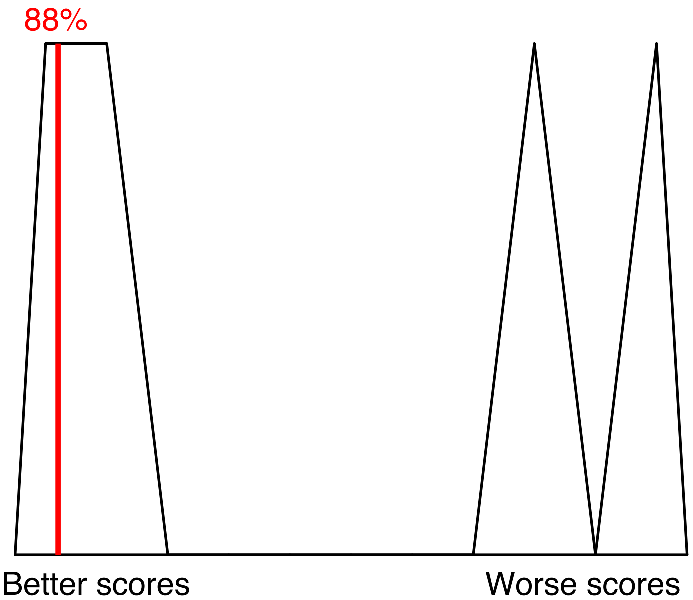

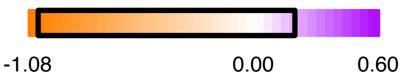

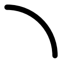
Protein Contact

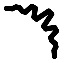
Protein Cavity

Residue Fingerprint

| | |
|---|---|
| ALA169A | ALA96A |
| ASP98A | GLU493A |
| GLY338A | GLY442A |
| ILE172A | LEU337A |
| LEU443A | PHE335A |
| PHE341A | SER336A |
| SER438A | SER439A |
| THR497A | TYR175A |
| TYR176A | TYR95A |

| | |
|---|---|
| Acceptor | Donor |
| Metal | Contact |

#### Shape -14.36

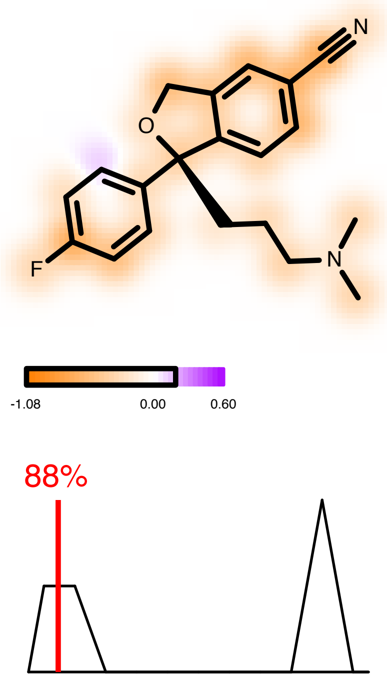

#### Hydrogen Bond 0.00

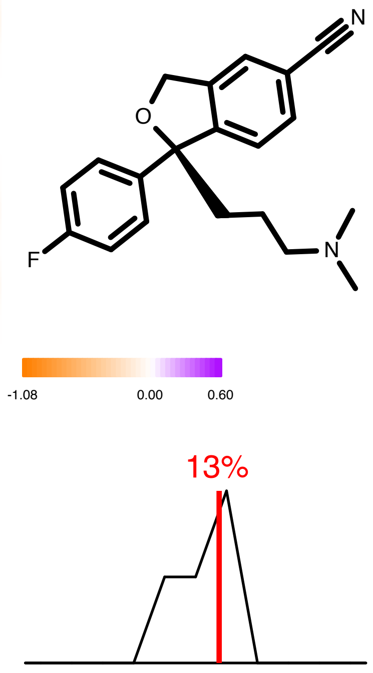

#### Protein Desolvation 2.50

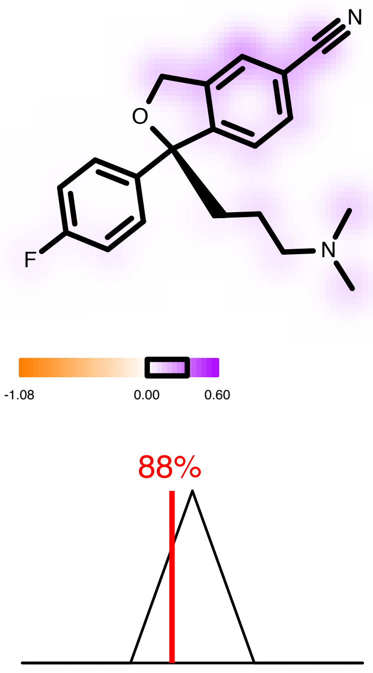

#### Ligand Desolvation 1.08

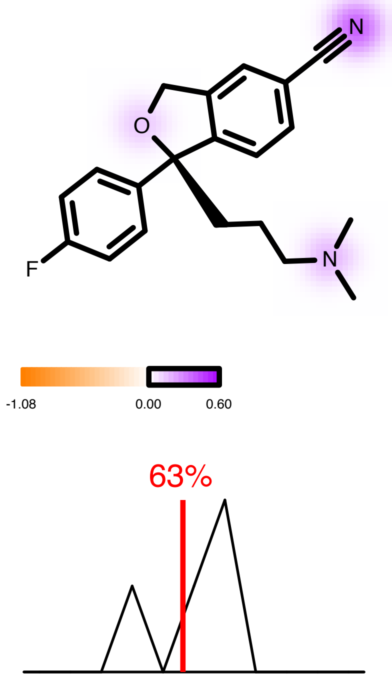

| | |
|---|---|
| Molecule Name | Fluvoxamine |
| Molecular Weight | 318.3 |
| XLogP | 4.1 |
| PSA | 56.8 |
| Heavy Atoms | 22 |
| Acceptor Count | 3 |
| Donor Count | 1 |
| Chelator Count | 0 |

### Total Score -10.44

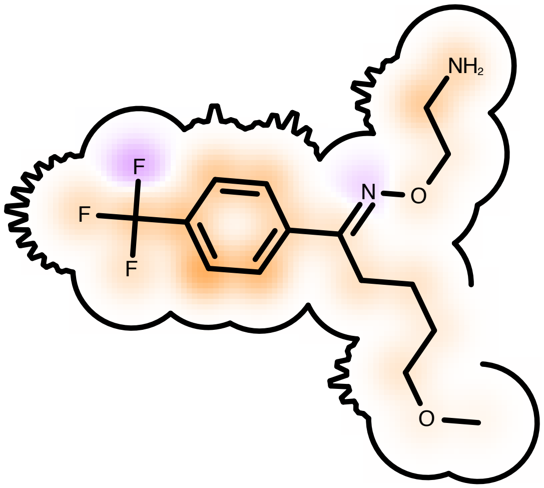

Score compared to other molecules

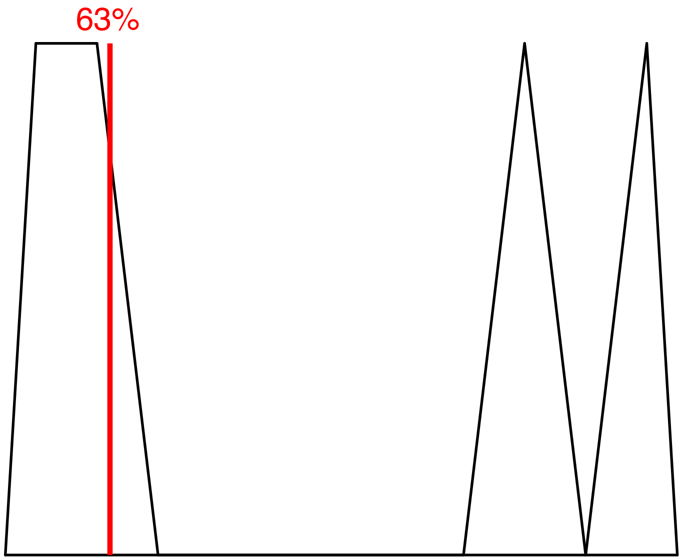

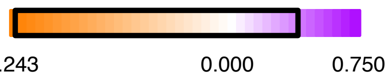

Protein Contact

Protein Cavity

<u>Residue Fingerprint</u>

| | |
|---|---|
| ALA169A | ALA96A |
| ASP98A | GLU493A |
| GLY338A | GLY442A |
| ILE172A | LEU337A |
| LEU443A | **PHE335A** |
| PHE341A | SER336A |
| SER438A | SER439A |
| THR497A | TYR175A |
| TYR176A | TYR95A |

**Acceptor** **Donor**
Metal Contact

#### Shape -14.04

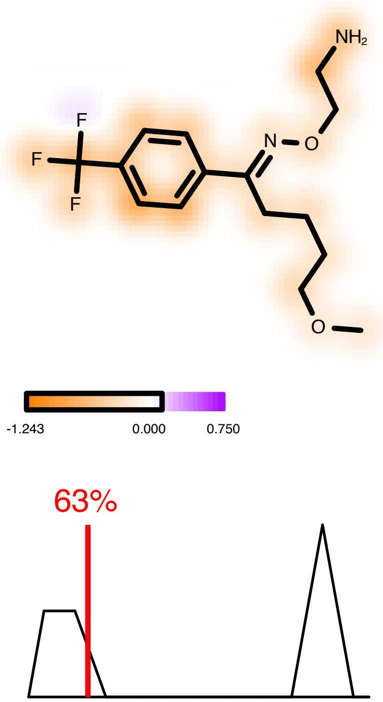

#### Hydrogen Bond -0.68

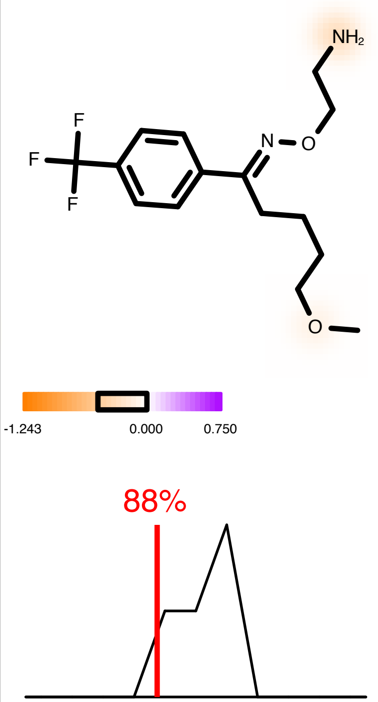

#### Protein Desolvation 2.63

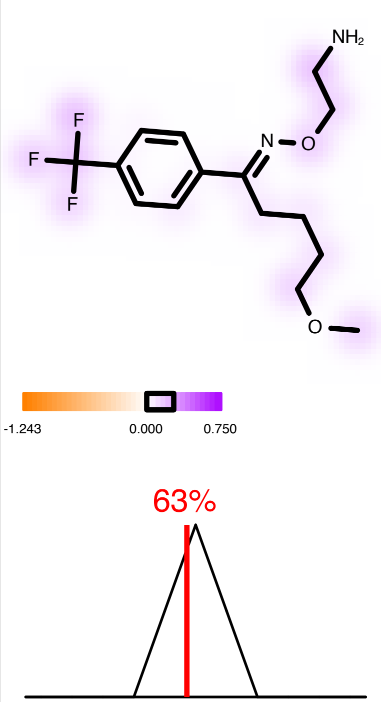

#### Ligand Desolvation 1.65

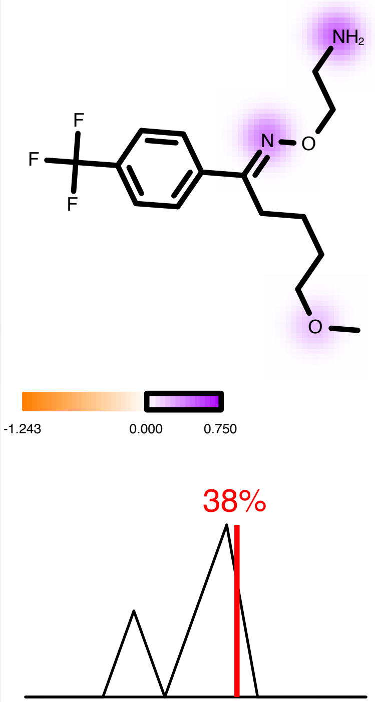

| | |
|---|---|
| Molecule Name | Sertraline |
| Molecular Weight | 306.2 |
| XLogP | 5.5 |
| PSA | 12.0 |
| Heavy Atoms | 20 |
| Acceptor Count | 1 |
| Donor Count | 1 |
| Chelator Count | 0 |

### Total Score -8.02

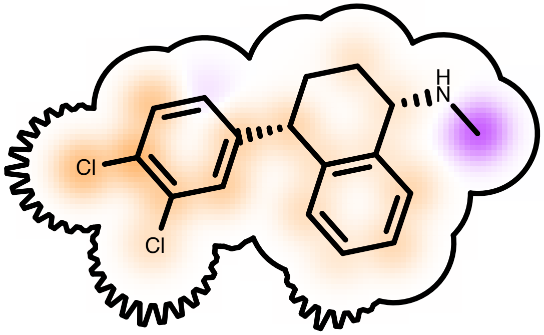

Score compared to other molecules

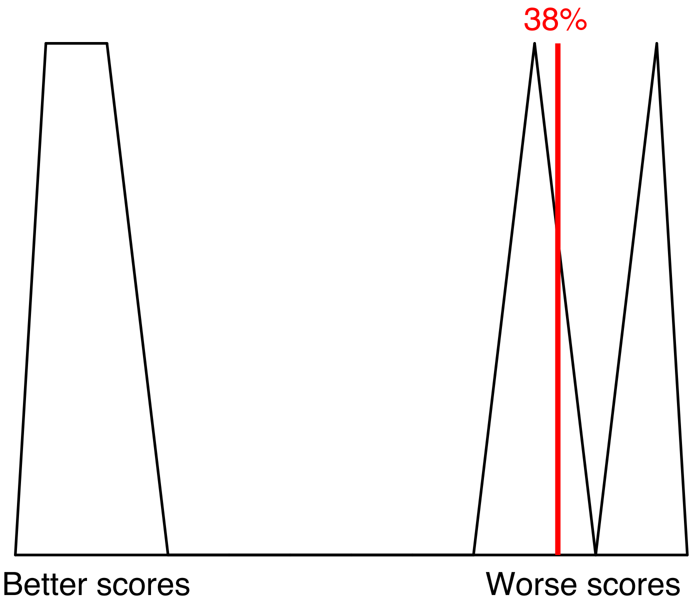

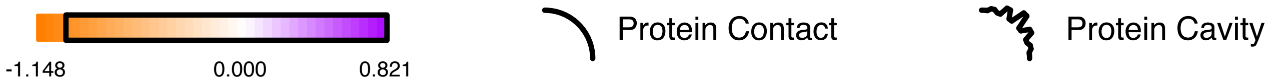

Residue Fingerprint

| | |
|---|---|
| ALA169A | ALA96A |
| ASP98A | GLU493A |
| GLY338A | GLY442A |
| ILE172A | LEU337A |
| LEU443A | PHE335A |
| PHE341A | SER336A |
| SER438A | SER439A |
| THR497A | TYR175A |
| TYR176A | TYR95A |

| | |
|---|---|
| Acceptor | Donor |
| Metal | Contact |

Shape -11.43

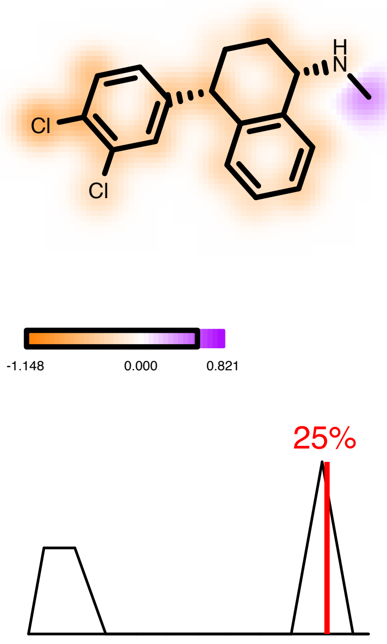

Hydrogen Bond -0.05

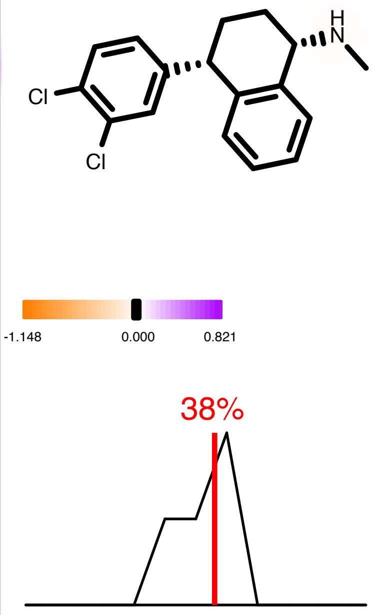

Protein Desolvation 3.04

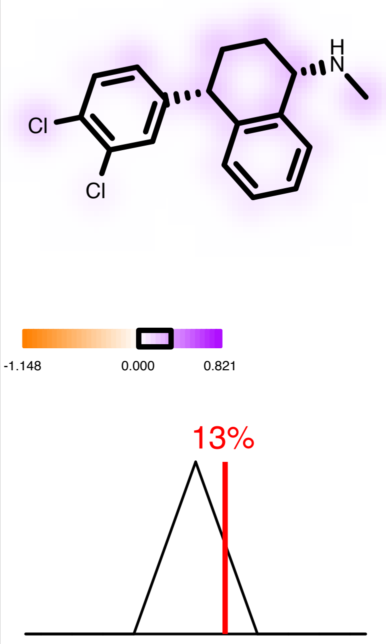

Ligand Desolvation 0.41

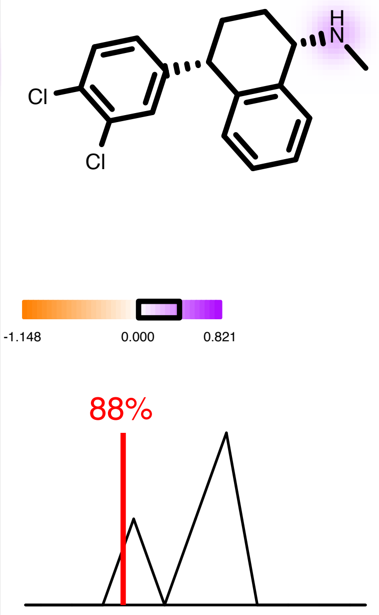

| | |
|---|---|
| Molecule Name | Paroxetine |
| Molecular Weight | 329.4 |
| XLogP | 3.4 |
| PSA | 39.7 |
| Heavy Atoms | 24 |
| Acceptor Count | 4 |
| Donor Count | 1 |
| Chelator Count | 0 |

### Total Score -7.41

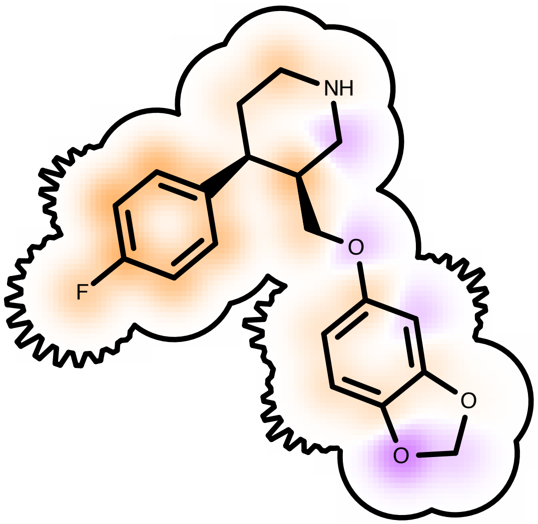

Score compared to other molecules

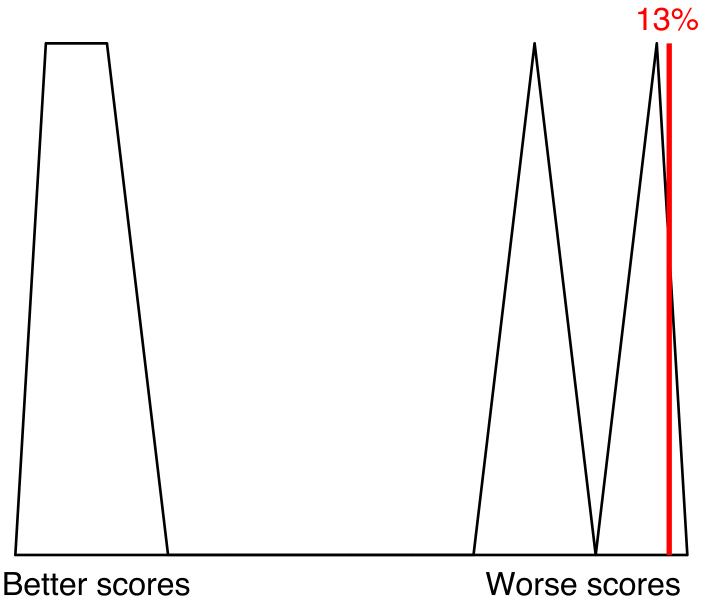

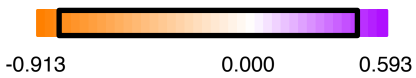

Protein Contact

Protein Cavity

Residue Fingerprint

| | |
|---|---|
| ALA169A | ALA96A |
| ASP98A | GLU493A |
| GLY338A | GLY442A |
| ILE172A | LEU337A |
| LEU443A | PHE335A |
| PHE341A | SER336A |
| SER438A | SER439A |
| THR497A | TYR175A |
| TYR176A | TYR95A |

| | |
|---|---|
| Acceptor | Donor |
| Metal | Contact |

#### Shape -11.49

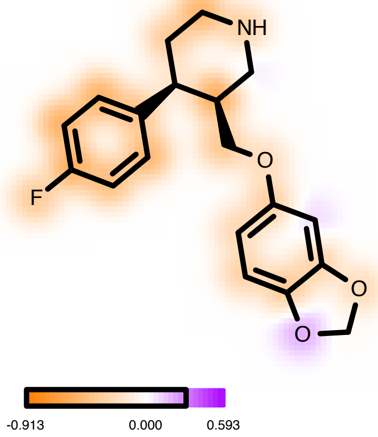

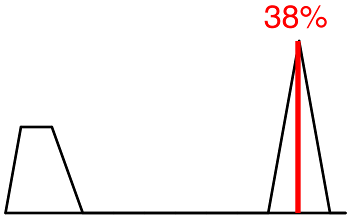

#### Hydrogen Bond -0.30

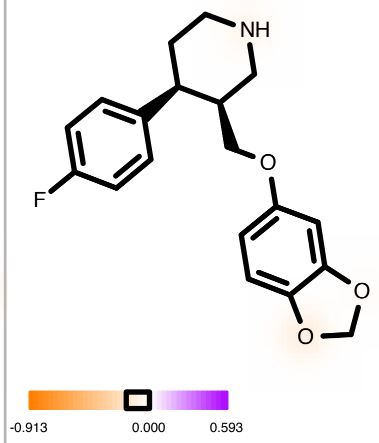

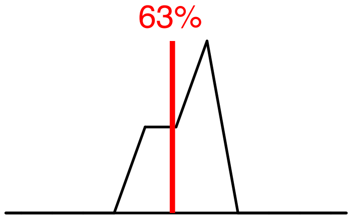

#### Protein Desolvation 2.72

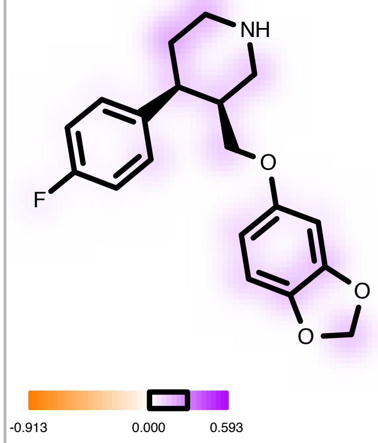

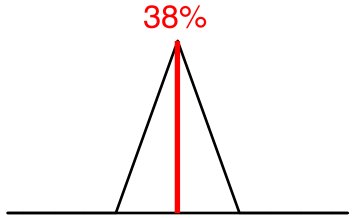

#### Ligand Desolvation 1.66

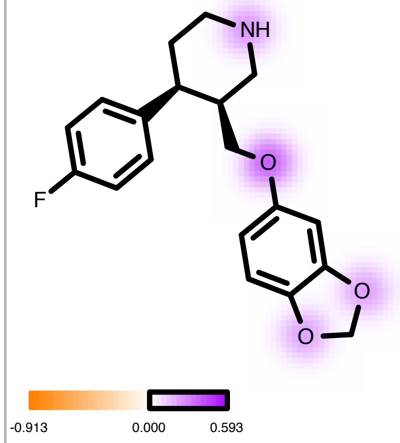

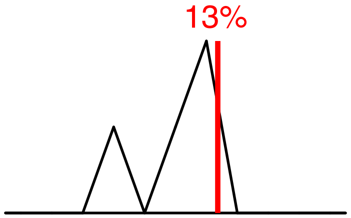

| | |
|---|---|
| Molecule Name | Drug Candidate 1 |
| Molecular Weight | 333.3 |
| XLogP | 5.3 |
| PSA | 20.2 |
| Heavy Atoms | 22 |
| Acceptor Count | 1 |
| Donor Count | 1 |
| Chelator Count | 1 |

### Total Score -11.88

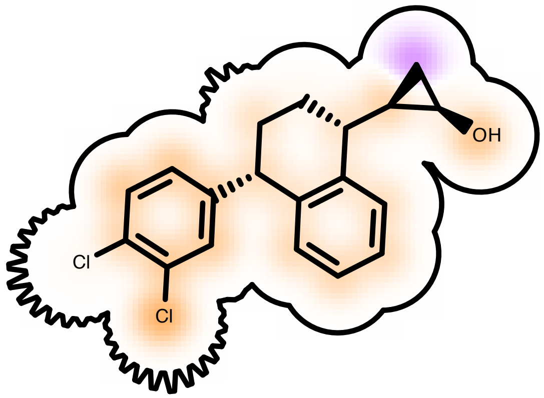

Score compared to other molecules

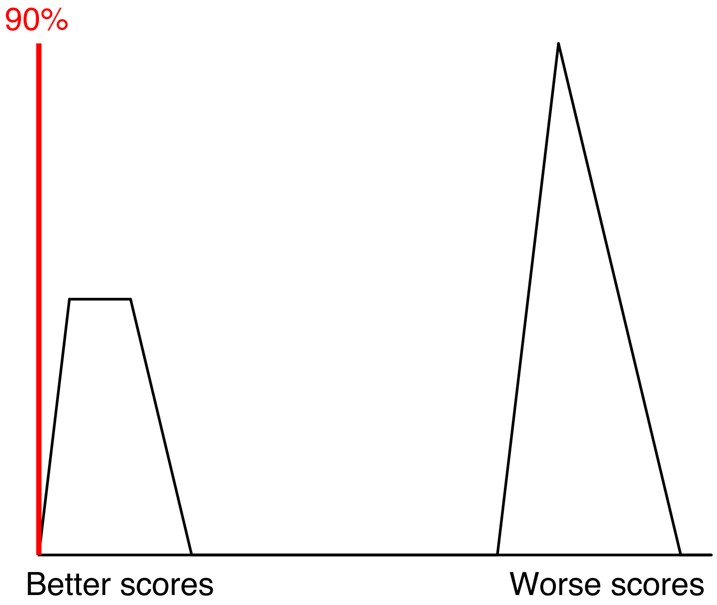

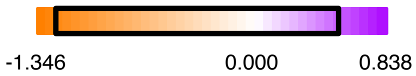

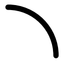
Protein Contact

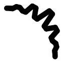
Protein Cavity

Residue Fingerprint

| | |
|---|---|
| ALA169A | ALA96A |
| ARG104A | ASN101A |
| ASP98A | GLY100A |
| GLY338A | GLY442A |
| ILE172A | LEU337A |
| PHE335A | PHE341A |
| SER336A | SER438A |
| SER439A | TYR176A |
| TYR95A | VAL343A |

| | |
|---|---|
| Acceptor | Donor |
| Metal | Contact |

#### Shape -14.59

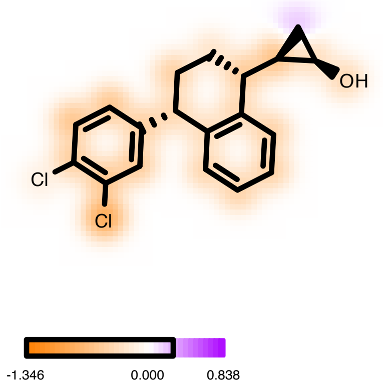

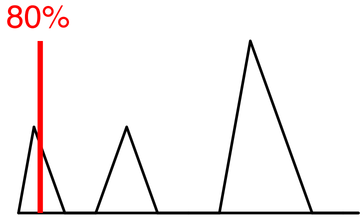

#### Hydrogen Bond -1.18

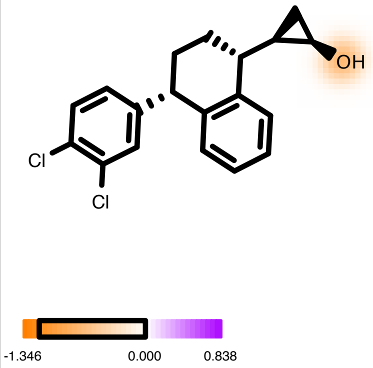

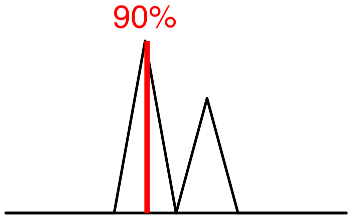

#### Protein Desolvation 3.05

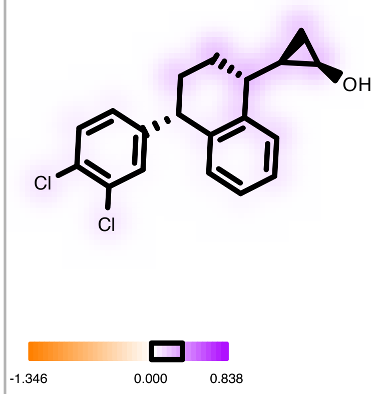

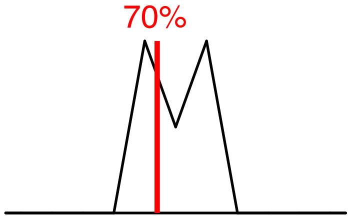

#### Ligand Desolvation 0.84

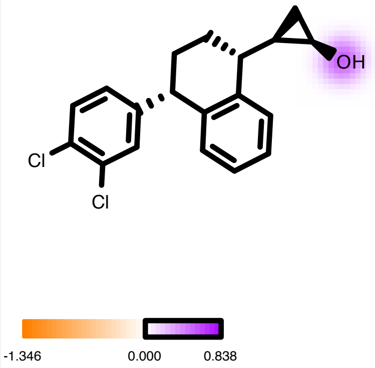

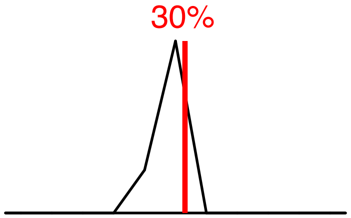

| | |
|---|---|
| Molecule Name | Drug Candidate 2 |
| Molecular Weight | 351.3 |
| XLogP | 4.8 |
| PSA | 29.5 |
| Heavy Atoms | 23 |
| Acceptor Count | 2 |
| Donor Count | 1 |
| Chelator Count | 1 |

### Total Score -11.56

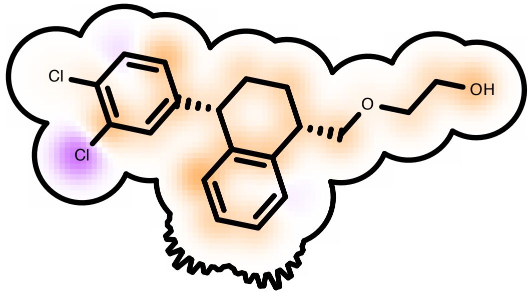

Score compared to other molecules

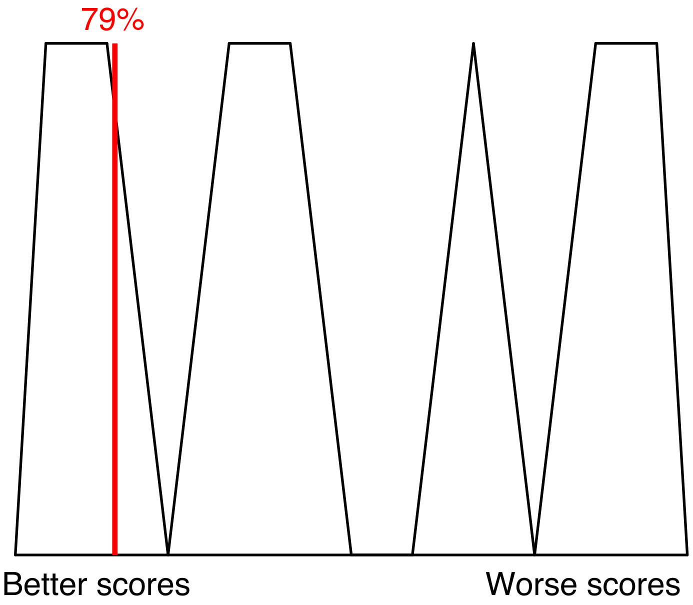

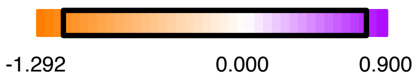

Protein Contact

Protein Cavity

Residue Fingerprint

| | |
|---|---|
| ALA169A | ALA173A |
| ALA96A | ARG104A |
| ASN101A | ASN177A |
| ASP98A | CYS473A |
| GLN332A | GLU493A |
| GLU494A | GLY100A |
| GLY338A | GLY442A |
| ILE172A | LEU337A |
| LEU443A | PHE263A |
| PHE335A | PHE341A |
| SER336A | SER438A |
| SER439A | THR497A |
| TYR175A | TYR176A |
| TYR95A | |

| | |
|---|---|
| Acceptor | Donor |
| Metal | Contact |

#### Shape -14.43

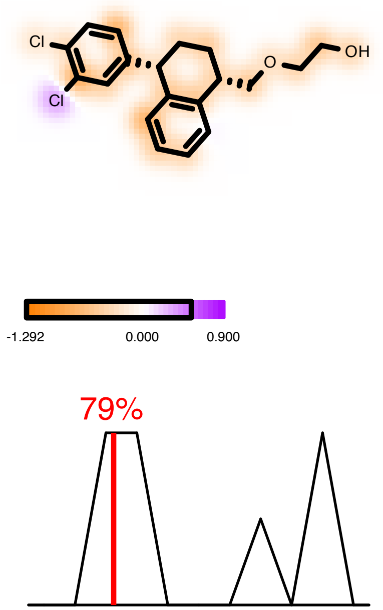

#### Hydrogen Bond -1.22

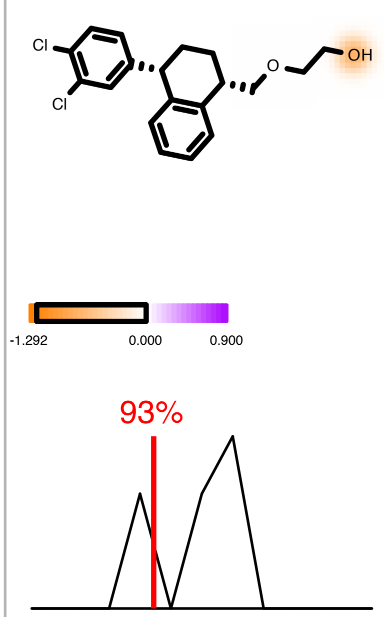

#### Protein Desolvation 2.88

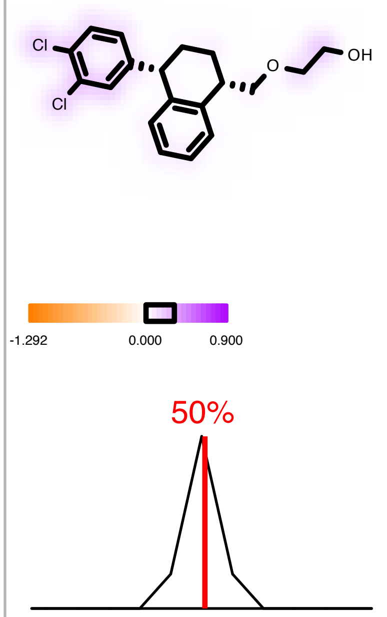

#### Ligand Desolvation 1.20

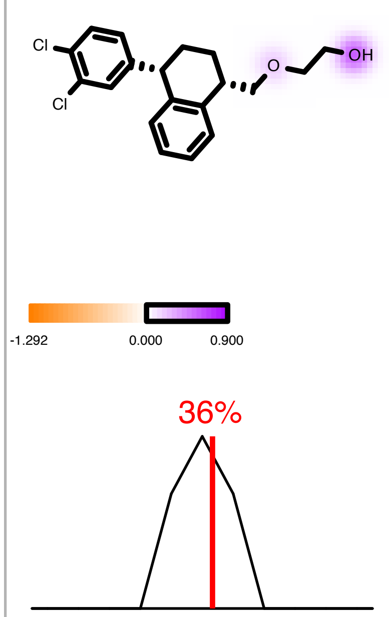
