## Supplemental Homology Analyses of the ts3 hSERT for "Design of Novel Selective Serotonin Reuptake Inhibitors Using Computational Modeling Studies"

**Supporting Information – Homology Analysis of the ts3 hSERT**

SciFinder searches of drug candidates 1 and 2 are listed in Fig. S-1 and Fig. S-2, respectively, whereas *ProteinBlast* homology analyses of clinical trial organisms and of beneficial and pathogenic microbes are listed in Fig. S-3, Fig. S-5, and Fig. S-6, respectively. Lastly, the conservation of Ser336 in the clinical model organisms, illustrated using *JalView*, is found in Fig. S-4.

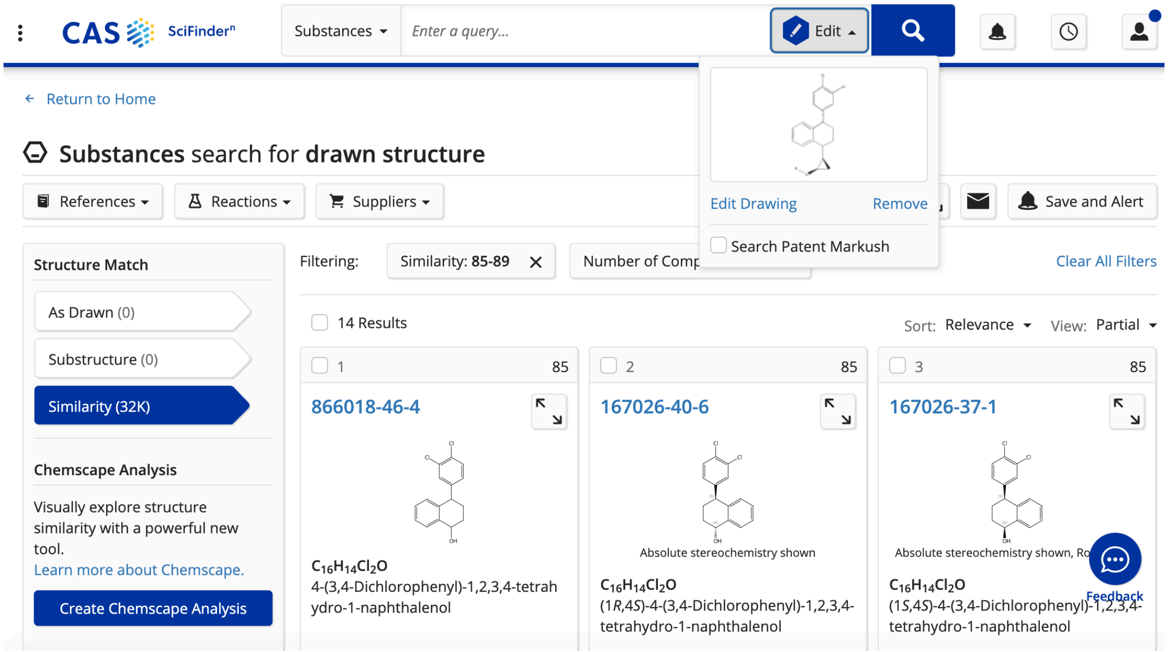

**Fig. S-1.** SciFinder search of drug candidate 1.

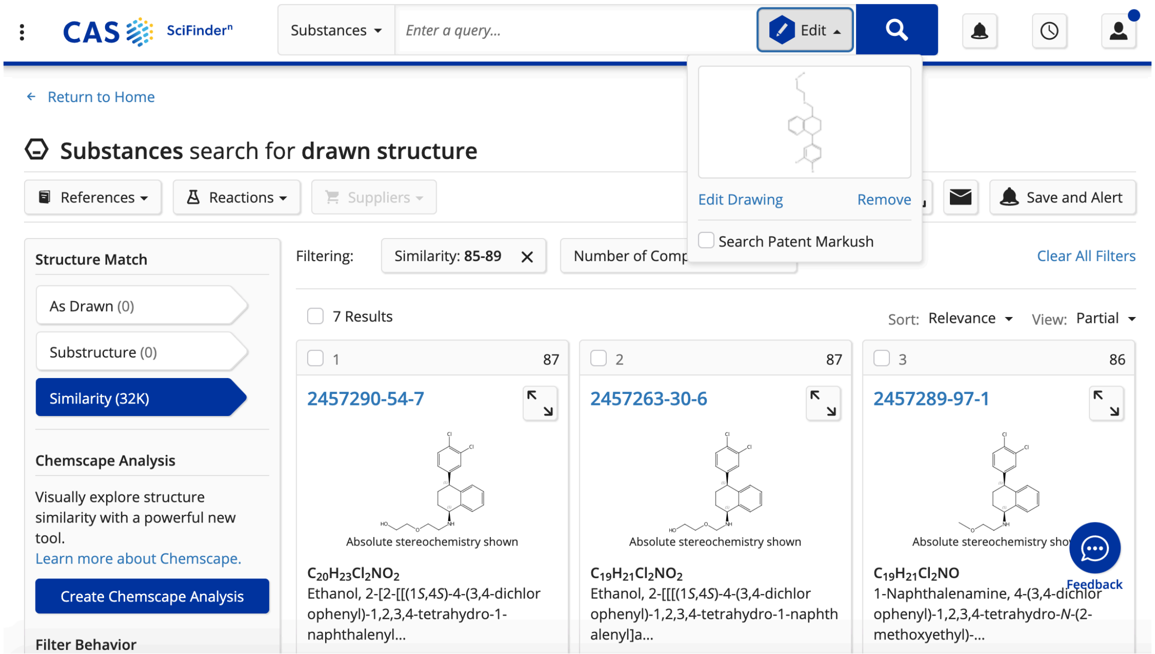

**Fig. S-2.** SciFinder search of drug candidate 2.

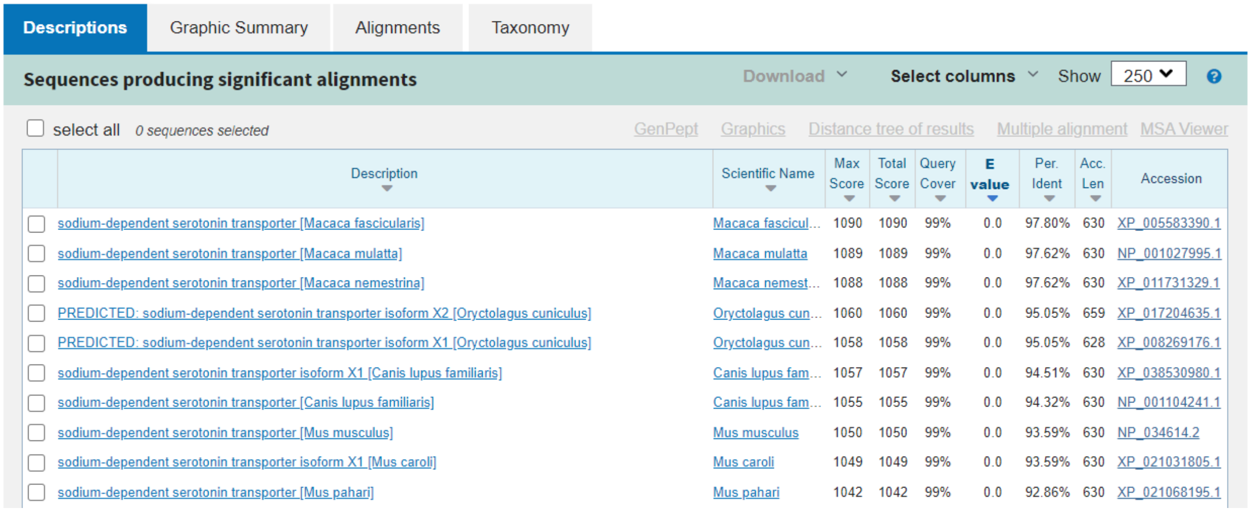

| Description | Scientific Name | Max Score | Total Score | Query Cover | E value | Per. Ident | Acc. Len | Accession |
|---|---|---|---|---|---|---|---|---|
| sodium-dependent serotonin transporter [Macaca fascicularis] | Macaca fascicul... | 1090 | 1090 | 99% | 0.0 | 97.80% | 630 | XP_005583390.1 |
| sodium-dependent serotonin transporter [Macaca mulatta] | Macaca mulatta | 1089 | 1089 | 99% | 0.0 | 97.62% | 630 | NP_001027995.1 |
| sodium-dependent serotonin transporter [Macaca nemestrina] | Macaca nemest... | 1088 | 1088 | 99% | 0.0 | 97.62% | 630 | XP_011731329.1 |
| PREDICTED: sodium-dependent serotonin transporter isoform X2 [Oryctolagus cuniculus] | Oryctolagus cun... | 1060 | 1060 | 99% | 0.0 | 95.05% | 659 | XP_017204635.1 |
| PREDICTED: sodium-dependent serotonin transporter isoform X1 [Oryctolagus cuniculus] | Oryctolagus cun... | 1058 | 1058 | 99% | 0.0 | 95.05% | 628 | XP_008269176.1 |
| sodium-dependent serotonin transporter isoform X1 [Canis lupus familiaris] | Canis lupus fam... | 1057 | 1057 | 99% | 0.0 | 94.51% | 630 | XP_038530980.1 |
| sodium-dependent serotonin transporter [Canis lupus familiaris] | Canis lupus fam... | 1055 | 1055 | 99% | 0.0 | 94.32% | 630 | NP_001104241.1 |
| sodium-dependent serotonin transporter [Mus musculus] | Mus musculus | 1050 | 1050 | 99% | 0.0 | 93.59% | 630 | NP_034614.2 |
| sodium-dependent serotonin transporter isoform X1 [Mus caroli] | Mus caroli | 1049 | 1049 | 99% | 0.0 | 93.59% | 630 | XP_021031805.1 |
| sodium-dependent serotonin transporter [Mus pahari] | Mus pahari | 1042 | 1042 | 99% | 0.0 | 92.86% | 630 | XP_021068195.1 |

**Fig. S-3.** Homology analysis of potential clinical trial organisms.

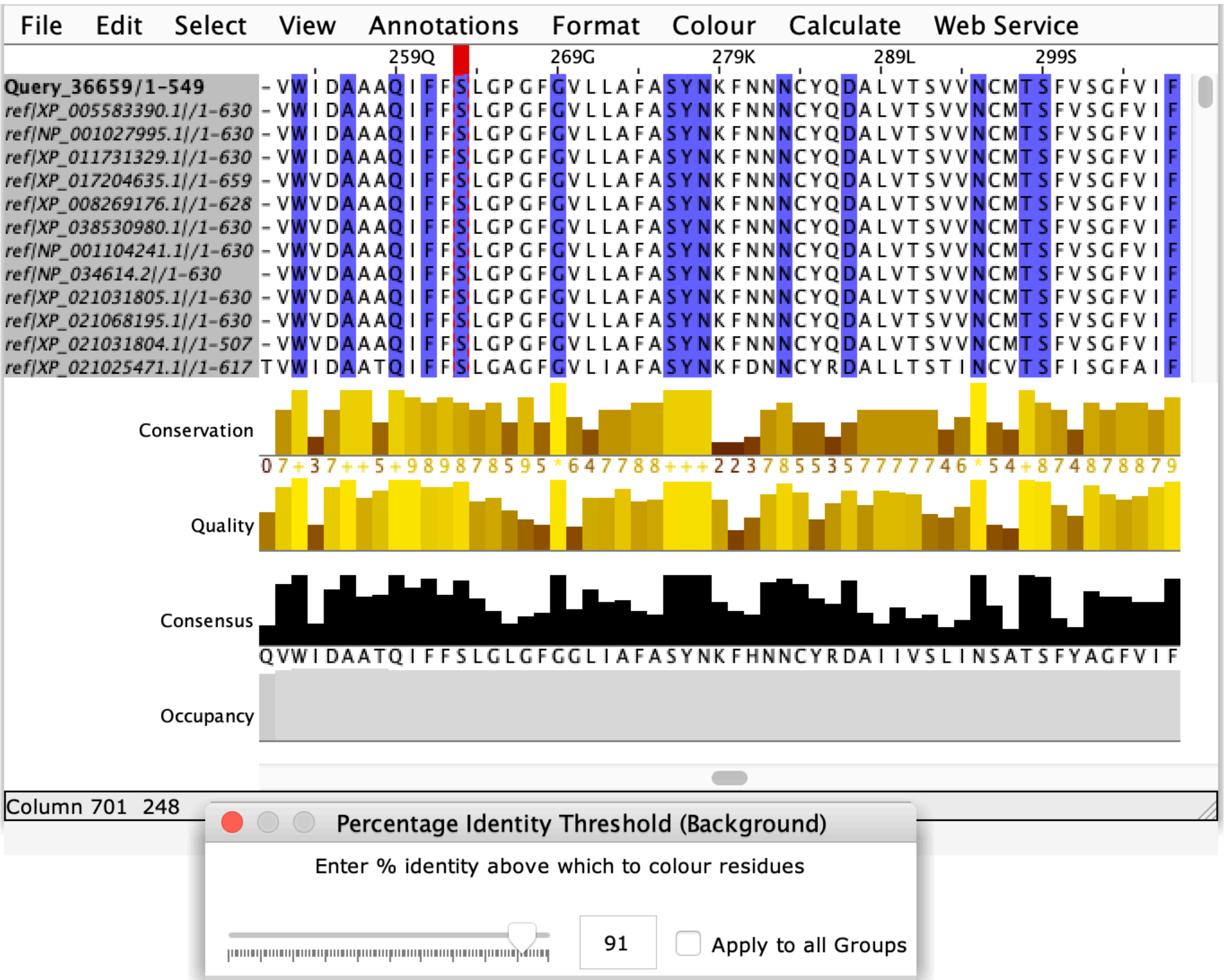

**Fig. S-4.** Conservation of Ser336 in clinical trial organisms.

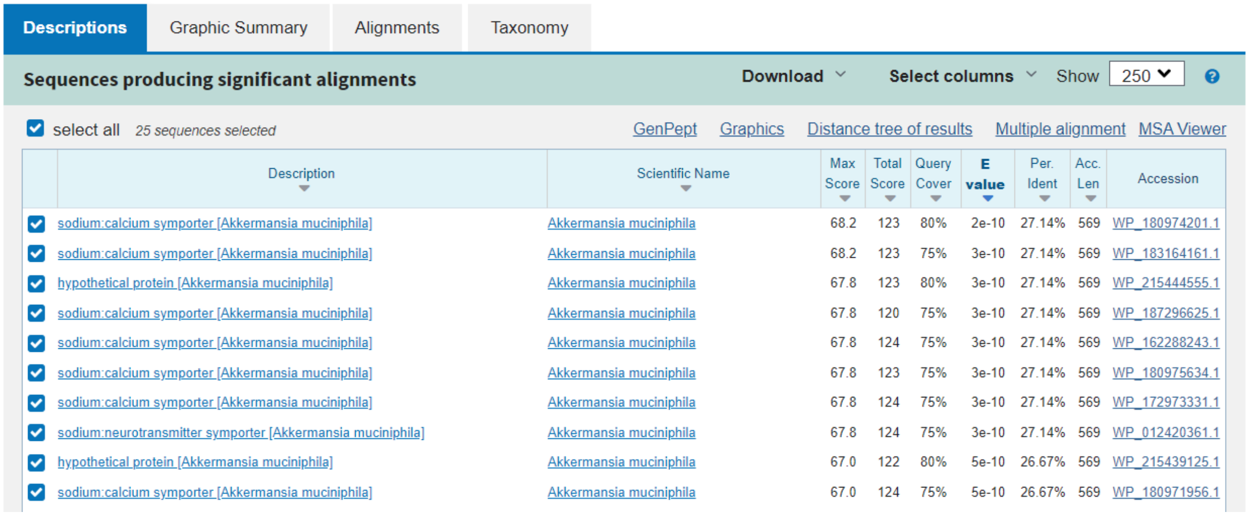

Descriptions | Graphic Summary | Alignments | Taxonomy

Sequences producing significant alignments — Download — Select columns — Show 250

select all 25 sequences selected — GenPept — Graphics — Distance tree of results — Multiple alignment — MSA Viewer

| Description | Scientific Name | Max Score | Total Score | Query Cover | E value | Per. Ident | Acc. Len | Accession |
|---|---|---|---|---|---|---|---|---|
| sodium:calcium symporter [Akkermansia muciniphila] | Akkermansia muciniphila | 68.2 | 123 | 80% | 2e-10 | 27.14% | 569 | WP_180974201.1 |
| sodium:calcium symporter [Akkermansia muciniphila] | Akkermansia muciniphila | 68.2 | 123 | 75% | 3e-10 | 27.14% | 569 | WP_183164161.1 |
| hypothetical protein [Akkermansia muciniphila] | Akkermansia muciniphila | 67.8 | 123 | 80% | 3e-10 | 27.14% | 569 | WP_215444555.1 |
| sodium:calcium symporter [Akkermansia muciniphila] | Akkermansia muciniphila | 67.8 | 120 | 75% | 3e-10 | 27.14% | 569 | WP_187296625.1 |
| sodium:calcium symporter [Akkermansia muciniphila] | Akkermansia muciniphila | 67.8 | 124 | 75% | 3e-10 | 27.14% | 569 | WP_162288243.1 |
| sodium:calcium symporter [Akkermansia muciniphila] | Akkermansia muciniphila | 67.8 | 123 | 75% | 3e-10 | 27.14% | 569 | WP_180975634.1 |
| sodium:calcium symporter [Akkermansia muciniphila] | Akkermansia muciniphila | 67.8 | 124 | 75% | 3e-10 | 27.14% | 569 | WP_172973331.1 |
| sodium:neurotransmitter symporter [Akkermansia muciniphila] | Akkermansia muciniphila | 67.8 | 124 | 75% | 3e-10 | 27.14% | 569 | WP_012420361.1 |
| hypothetical protein [Akkermansia muciniphila] | Akkermansia muciniphila | 67.0 | 122 | 80% | 5e-10 | 26.67% | 569 | WP_215439125.1 |
| sodium:calcium symporter [Akkermansia muciniphila] | Akkermansia muciniphila | 67.0 | 124 | 75% | 5e-10 | 26.67% | 569 | WP_180971956.1 |

**Fig. S-5.** Homology analysis of beneficial microbes.

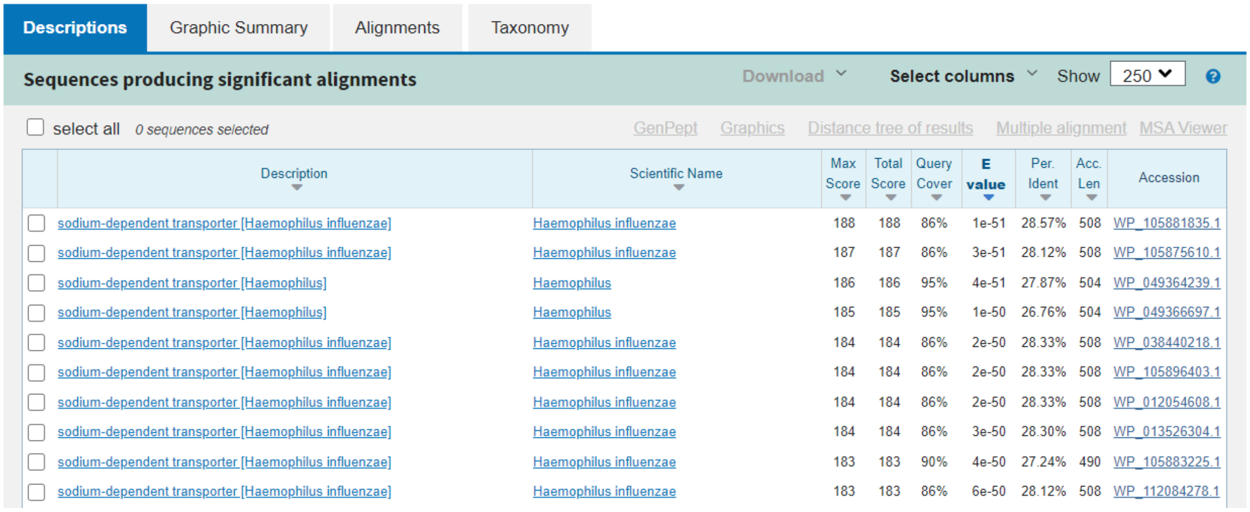

Descriptions | Graphic Summary | Alignments | Taxonomy

Sequences producing significant alignments — Download — Select columns — Show 250

select all 0 sequences selected — GenPept — Graphics — Distance tree of results — Multiple alignment — MSA Viewer

| Description | Scientific Name | Max Score | Total Score | Query Cover | E value | Per. Ident | Acc. Len | Accession |
|---|---|---|---|---|---|---|---|---|
| sodium-dependent transporter [Haemophilus influenzae] | Haemophilus influenzae | 188 | 188 | 86% | 1e-51 | 28.57% | 508 | WP_105881835.1 |
| sodium-dependent transporter [Haemophilus influenzae] | Haemophilus influenzae | 187 | 187 | 86% | 3e-51 | 28.12% | 508 | WP_105875610.1 |
| sodium-dependent transporter [Haemophilus] | Haemophilus | 186 | 186 | 95% | 4e-51 | 27.87% | 504 | WP_049364239.1 |
| sodium-dependent transporter [Haemophilus] | Haemophilus | 185 | 185 | 95% | 1e-50 | 26.76% | 504 | WP_049366697.1 |
| sodium-dependent transporter [Haemophilus influenzae] | Haemophilus influenzae | 184 | 184 | 86% | 2e-50 | 28.33% | 508 | WP_038440218.1 |
| sodium-dependent transporter [Haemophilus influenzae] | Haemophilus influenzae | 184 | 184 | 86% | 2e-50 | 28.33% | 508 | WP_105896403.1 |
| sodium-dependent transporter [Haemophilus influenzae] | Haemophilus influenzae | 184 | 184 | 86% | 2e-50 | 28.33% | 508 | WP_012054608.1 |
| sodium-dependent transporter [Haemophilus influenzae] | Haemophilus influenzae | 184 | 184 | 86% | 3e-50 | 28.30% | 508 | WP_013526304.1 |
| sodium-dependent transporter [Haemophilus influenzae] | Haemophilus influenzae | 183 | 183 | 90% | 4e-50 | 27.24% | 490 | WP_105883225.1 |
| sodium-dependent transporter [Haemophilus influenzae] | Haemophilus influenzae | 183 | 183 | 86% | 6e-50 | 28.12% | 508 | WP_112084278.1 |

**Fig. S-6.** Homology analysis of pathogenic microbes.
